# A genome-wide *in vivo* screen reveals fitness pathways required for streptococcal infective endocarditis

**DOI:** 10.64898/2026.04.07.717129

**Authors:** Liang Bao, Jennifer Bradley, Vysakh Anandan, Katarzyna M. Tyc, Zan Zhu, Josephina Anna Vossen, Valery-francine Assi, Jasmine S. Benbei, Nicai Zollar, Todd Kitten, Ping Xu

## Abstract

Infective endocarditis (IE) is a life-threatening disease most often caused by blood-borne bacteria that infect previously damaged cardiac tissue. Despite the importance of this disease, the genetic basis for IE virulence remains poorly defined. Here, we present the first genome-wide *in vivo* analysis of bacterial fitness in a vertebrate model of IE. We identified 146 genes in *Streptococcus sanguinis* required for IE fitness, the majority of which had not previously been linked to endocarditis. These determinants cluster into conserved metabolic, cell envelope, transport, and regulatory pathways, representing a vast reservoir of potential targets for novel antimicrobial intervention. A subset of these genes was examined in *Streptococcus mutans*; all were found to be essential for IE fitness in this distantly related oral species as well, suggesting broad conservation. Using experimental evolution, we further show that disruption of key fitness pathways triggers reproducible compensatory “bypass” mechanisms that reveal the inherent physiological constraints of the IE fitness landscape and identify vulnerable nodes for multi-target drug strategies. Together, these findings redefine streptococcal infective endocarditis as a disease shaped by conserved bacterial fitness networks that may be exploited for therapeutic development.

**Highlights:**

- A genome-wide *in vivo* screen identified 146 *Streptococcus sanguinis* genes required for infective endocarditis fitness.
- 94% of these genes represent previously unrecognized determinants of endocarditis.
- Multiple pathways—including CoA biosynthesis, the shikimate pathway, and rhamnan synthesis—are required for cardiac colonization.
- A subset of infective endocarditis fitness factors are conserved between *S. sanguinis* and *S. mutans*, with species-specific adaptations.
- Experimental evolution revealed compensatory metabolic networks that buffer IE fitness defects.

## Introduction

Infective endocarditis (IE) is a life-threatening infection of the endocardial surface of the heart, frequently leading to severe complications such as valvular destruction, systemic embolization, and heart failure[1, 2]. Despite advancements in medical care[2, 3], the mortality associated with IE remains alarmingly high, with in-hospital mortality rates ranging from 15% to 20% and a 1-year mortality rate approaching 40%[4]. The pathogenesis of IE is driven by complex interactions between the bacteria and the host[1, 2]. Typically, streptococcal IE begins with a lesion or minor defect in the endocardial endothelium, which induces the formation of a sterile, healing vegetation composed of fibrin and platelets[5–7]. This vegetation provides a nidus for bacterial colonization during transient bacteremia. This is followed by biofilm-like bacterial growth, enlargement of vegetative lesions, and inflammation of the tissues[5–7]. Thus, streptococcal IE progression is likely driven not by acute toxin-mediated damage but by the sustained capacity of IE pathogens to survive, grow, and persist within cardiac vegetations—an environment defined by nutrient limitation, immune pressure, and mechanical stress. However, the genome-wide genetic determinants that confer fitness in this specialized host niche remain poorly understood.

Oral streptococci are genetically diverse and differ in their contributions to oral and systemic diseases. These bacteria can enter the bloodstream during invasive dental procedures or routine activities such as toothbrushing or chewing, potentially leading to IE[8–10]. The prevention and management of IE are critical concerns, especially for patients with cardiac conditions that place them at elevated risk[10, 11]. While high-dose antibiotic prophylaxis is often recommended for these patients prior to invasive dental procedures, the protection afforded[12, 13] comes at the cost of increased antibiotic resistance in healthy individuals[14] and eradication of beneficial flora from the oral cavity. Moreover, it provides no protection from daily bacteremia[15–17]. The identification of targets whose inactivation prevents IE without affecting oral colonization could afford the benefits of standard prophylaxis without the attendant costs.

By employing animal models in which a catheter is introduced into the heart to create sterile cardiac vegetations prior to intravenous inoculation of bacteria[7, 18], we and others have identified a number of *S. sanguinis* fitness factors for IE[19–27]. These studies have been useful, but most have examined only a small number of genes for their impact on IE. Even the two studies that set out to examine every putative lipoprotein[20] or cell wall-anchored protein[28] for their contribution to IE examined only small subsets of the genes contained within the 2.4-Mbp *S. sanguinis* strain SK36 genome.

Genome-wide *in vivo* screening approaches potentially allow for comprehensive assessment of the bacterial genes required for disease causation in animal models. However, these methods face inherent challenges, particularly when testing mutant pools. Bottleneck effects, where only a random subset of mutants successfully colonizes infection sites, can skew fitness outcomes and lead to inconsistent results[29]. Additionally, fitness levels of specific mutants, generally evaluated using competitiveness within a pool, often vary across experiments, further complicating the identification of consistently attenuated mutants. Indeed, these issues limited the interpretability of a previous pooled mutant screen in *S. sanguinis* IE[19]. Moreover, fitness screens alone do not reveal how disrupted pathways support bacterial physiology or how bacteria adapt when these pathways are compromised, leaving the regulatory and compensatory logic of IE fitness largely unexplored.

Here, we address these challenges by combining a comprehensive nonessential gene deletion library[30] in *S. sanguinis* with a next-generation sequencing–based ORFseq platform[31] and a rigorously optimized vertebrate IE model[7, 18]. By conducting thousands of competitive fitness measurements across independent animal experiments, we systematically assessed the contribution of nearly every annotated open reading frame to IE-associated fitness. This approach enabled us to identify high-confidence fitness determinants, define conserved and species-specific IE fitness pathways, and uncover potential adaptive compensatory mechanisms through experimental evolution. Together, our findings provide a genome-wide framework for understanding the physiological basis of IE virulence and reveal conserved bacterial vulnerabilities.

## Results

### *In vivo* screening for identification of fitness factors

To systematically investigate the genetic basis of *Streptococcus sanguinis* fitness in IE, we performed a genome-wide screen to identify gene deletions that influence competitiveness of pooled mutants in our *in vivo* model. The model employs introduction of a catheter into the heart to stimulate the formation of sterile vegetations, followed by inoculation of bacteria through an ear vein (see Materials and Methods) (Figure 1A). The mutants tested were derived from our comprehensive non-essential gene knockout library, comprising 2,011 mutants[30], 27 newly annotated open reading frame (ORF) knockout mutants, and one essential gene mutant, Δ*f1fo* [32], for a total of 2,039 mutants (Table S1). These mutants were tested in 24 pools, with sizes of 85 to 212 mutants per pool (Figure 1A and 1B; Table S2).

**Figure 1.**
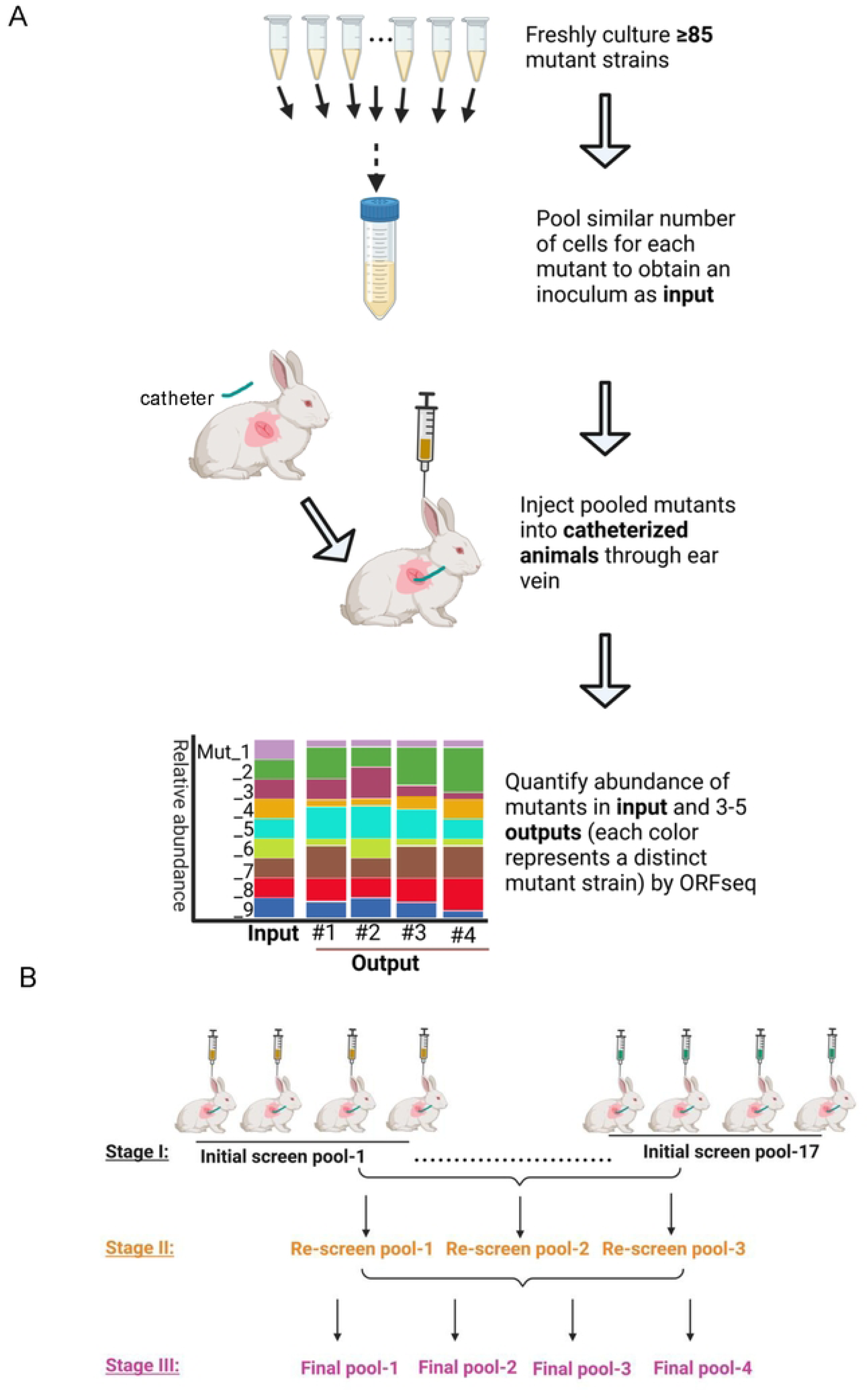
Experimental design and screening workflow for identification of IE-related fitness factors in *S. sangunis* SK36. **A.** Sequential steps for evaluating the fitness of mutants using ORF seq with the animal IE model (see Materials and Methods for detailed procedures). **B.** Overview of the stages of mutant screening, including initial screening, re-screening, and confirmation, conducted across a total of24 anin1al experiments.

To ensure the reliability of our screens, we included three known IE fitness-reduced mutants as positive controls—Δ*purB* (SSA_0046), Δ*ssaB* (SSA_0260), and Δ*ssaC* (SSA_0261)—and a hypothetical-protein mutant with wild-type-like fitness, ΔSSA_0169 (Table S3) as the negative control. Each pool was tested in three to five catheterized rabbits (Figure 1A; Table S3).

Fitness of mutants was assessed by calculating the ratio of mutant abundance in cardiac vegetations[21, 24] of catheterized animals approximately 20 hours post-inoculation relative to their abundance in the input pools—a measure we will refer to as the “abundance ratio.” In streptococcal endocarditis, pathology results primarily from growth of the bacteria and the resulting growth of the vegetation[7]; thus, *in vivo* growth is tantamount to IE virulence. Mutant abundance in input pools (the bacterial inocula for animals) and output vegetations was quantified using ORFseq[31], which identifies mutants via ORF-linked *APH(3’)-IIIa* (kanamycin resistance gene) tags (see Materials and Methods; Fig. S1).

### Pool Size Optimization and Validation

Our previous signature-tagged mutagenesis study[19] employed a pool size of 40 mutants using a similar rabbit model. To enhance screening efficiency without inducing bottleneck effects[33], we pilot tested 85, 212, and 159 mutants in Initial-Screen Pool-1, Pool-2, and Pool-3, respectively [33] (Tables S2 and S3). To reduce variation, we pooled similar cell numbers of each mutant in each experiment (Figure 1A; Fig. S1; see Materials and Methods).

For Pool 1, all three fitness-reduced controls (Δ*purB*, Δ*ssaB*, and Δ*ssaC*) exhibited reduced recovery, while the WT-like control (ΔSSA_0169)[34] was reproducibly recovered from every animal, confirming the pool’s validity. However, in Pool 2 (212 mutants), the WT control showed significantly reduced abundance, indicating a bottleneck effect[33]. We then examined Pool 3 (159 mutants), where fitness-reduced controls were confirmed, and the WT control was reproducibly recovered. Based on these results, we selected 159 mutants as the optimal pool size and employed this approximate pool size for the 21 remaining pools (Fig. S 2A-B; Tables S2, S3 and S4). All (100%) of the 66 tests of the fitness-reduced control mutants demonstrated significantly reduced abundance ratios, while 21 of 22 tests (95.45%) showed no significant reduction in abundance ratios for the WT control. The lone exception was “initial screen pool-2,” as discussed above.

### Identification, description, and conservation of 146 identified IE fitness determinants

During screening, the distribution of mutant fitness values revealed that although some mutants occasionally exhibited increased abundance in individual animals, these effects were inconsistent and likely reflected stochastic variation. Accordingly, we focused on mutants that consistently showed reduced fitness across replicate experiments, defined by an abundance ratio < 1 and statistically significant attenuation based on combined p-values and the number of independent tests (see Materials and Methods). Using these criteria, we identified 146 high-confidence fitness determinants (with the F₁F₀ ATPase operon treated as a single locus) (Tables S5–S6). Interestingly, 137 (94%) of the 146 candidates identified in this study are novel IE fitness factors in that they have not been previously associated with IE in any bacterium (Table S6). These include 17 that are annotated as hypothetical proteins with unknown functions.

To define functional relationships among the 146 IE fitness determinants identified in *S. sanguinis* SK36, these genes were grouped into five functional categories: DNA replication and cell division; transcription, translation and post-translational modification; cell wall synthesis; transport and metabolism; and hypothetical proteins (Figure 2). Genes involved in DNA replication and cell division encoded multiple purine biosynthetic enzymes and factors associated with DNA protection, primosome function, replication initiation, and septation. The transcription, translation, and post-translational modification category encompassed genes involved in transcriptional regulation, RNA degradation, ribosomal proteins, ribosome biogenesis, protein synthesis, proteolysis, and protein secretion. Cell wall synthesis genes included the *rml* and *rgp* genes required for rhamnan biosynthesis. Transport-related IE fitness determinants comprised core phosphotransferase system components (HPr and enzyme I), the metal transporter SsaACB, multiple energy-coupling factor (ECF) transporters, and the F1Fo ATPase. Metabolic genes were enriched for the shikimate pathway, CoA synthesis, pyridoxal phosphate–dependent aminotransferases, serine and tryptophan biosynthesis, and glycolytic enzymes. Seventeen IE fitness determinants were annotated as hypothetical proteins. In addition, two genes—SSA_1509, located within the *rgp* gene cluster, and SSA_2367, located within an ECF transporter operon—were not directly tested due to their absence from the input library but are likely IE fitness determinants based on genomic context.

**Figure 2.**
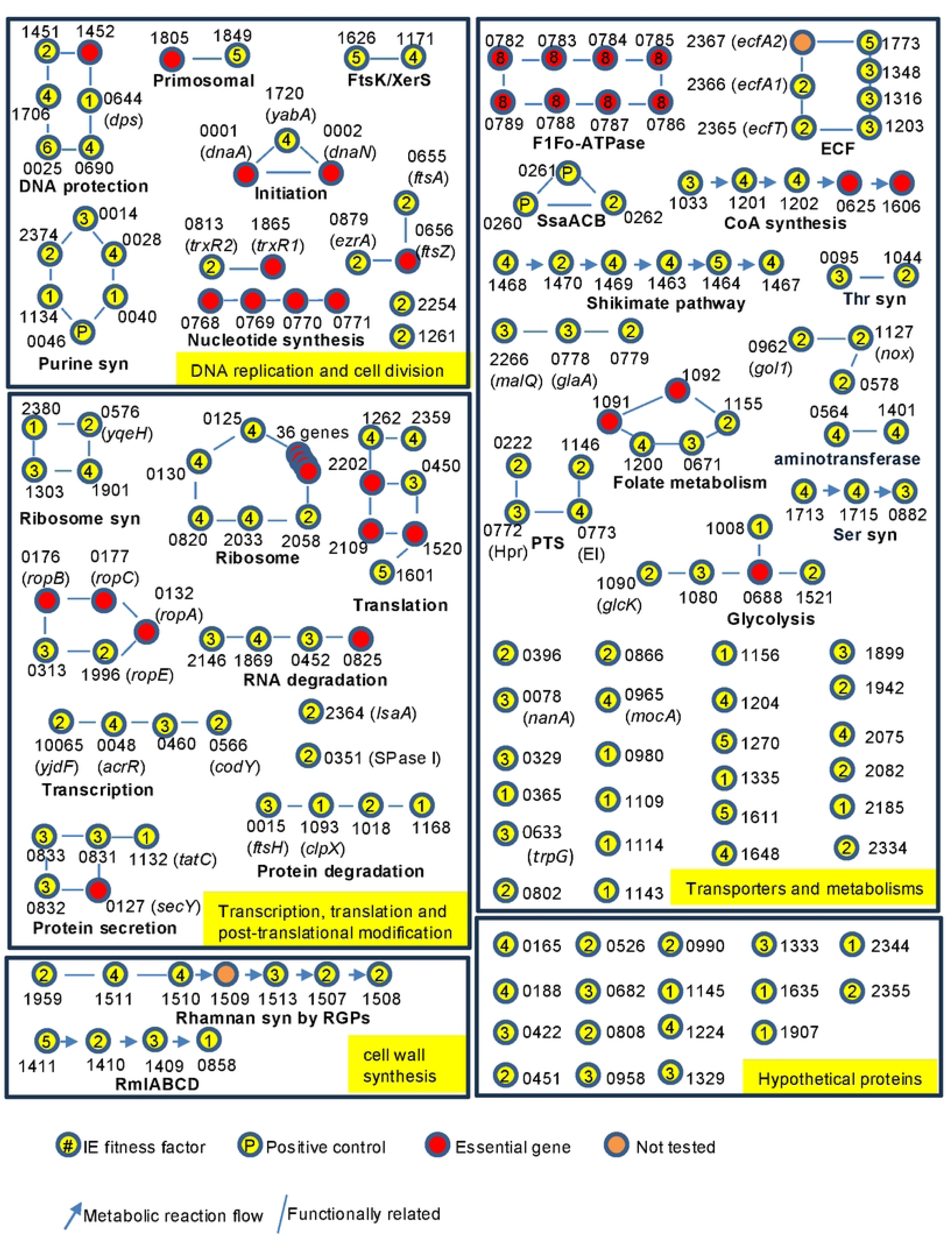
Interactions of IE fitness genes. The panel illustrates pathways in which IE fitness genes interact with other IE fitness genes, organized into five functional groups. Essential genes within each pathway are included. Pathway names are shown in bold beneath each network. Gene locus identifiers with four digits correspond to **SSA_XXXX,** whereas those with six digits correspond to **JlC87 XXXXX** Nu1nbers inside the circles indicate the number of independent fitness assays pe1fonned for each mutant.

Several IE fitness genes functionally intersected with pathways containing essential genes, highlighting their integration into core cellular processes. For example, the products of the IE fitness determinants *ftsA* (SSA_0655) and *ezrA* (SSA_0879) interact with the essential cell division protein FtsZ[35]; five IE fitness ribosomal proteins interface with an additional 36 essential ribosomal proteins; and the IE fitness genes *coaA*, *coaB*, and *coaC* act together with essential *coaD* and *coaE* in coenzyme A biosynthesis. In contrast, other IE fitness pathways—such as rhamnan biosynthesis (*rml* and *rgp* genes), purine biosynthesis, the SsaACB metal transporter, and the shikimate pathway—are composed largely of nonessential genes, suggesting requirements for infection that do not apply to *in vitro* growth.

To assess the evolutionary conservation of these IE fitness determinants, homology searches were performed across 194 genomes from the three genera of Gram-positive cocci that cause the majority of IE cases: streptococci; enterococci; and staphylococci (Tables S6 and S7). Comparative genomic analysis revealed extensive conservation of IE fitness genes within the genus *Streptococcus* and substantial conservation across genera. Most IE fitness determinants were present in nearly all *Streptococcus* strains and were enriched for functions related to DNA replication and repair, ribosome biogenesis and translation, central carbon metabolism, and energy homeostasis, indicating a requirement to maintain core physiological functions in the endocardial niche. Genes involved in amino acid, nucleotide, and cofactor biosynthesis—including the shikimate pathway, purine biosynthesis, folate-mediated one-carbon metabolism, and CoA synthesis—were also conserved. Additionally, genes involved in cell division, envelope integrity, and stress response were among the most conserved IE fitness determinants. While many IE fitness genes were shared across *Streptococcus*, *Enterococcus*, and *Staphylococcus*, a subset—including components of the accessory Sec secretion system, select glycosyltransferases, and several hypothetical or domain-of-unknown-function proteins—was present only within *Streptococcus*. Collectively, these data define a conserved Gram-positive IE fitness architecture composed of core metabolic and regulatory functions integrated with species-specific accessory systems.

We next examined whether IE fitness determinants clustered within shared metabolic pathways or protein complexes. Using genomic organization and functional annotation, we identified seven systems in which multiple genes contributed to IE fitness (Figure 3A): the shikimate pathway (Fig. S3A), CoA biosynthesis (Fig. S3B), rhamnan synthesis (*rlmABCD* and *rgpABCDF*; Fig. S3C), the EI and HPr components of the phosphotransferase system (PTS; Fig. S3D), energy-coupling factor (ECF) transporters (Fig. S3E), serine biosynthesis (Fig. S3F), and the SsaACB manganese transporter (Fig. S3G) [21, 36].

**Figure 3.**
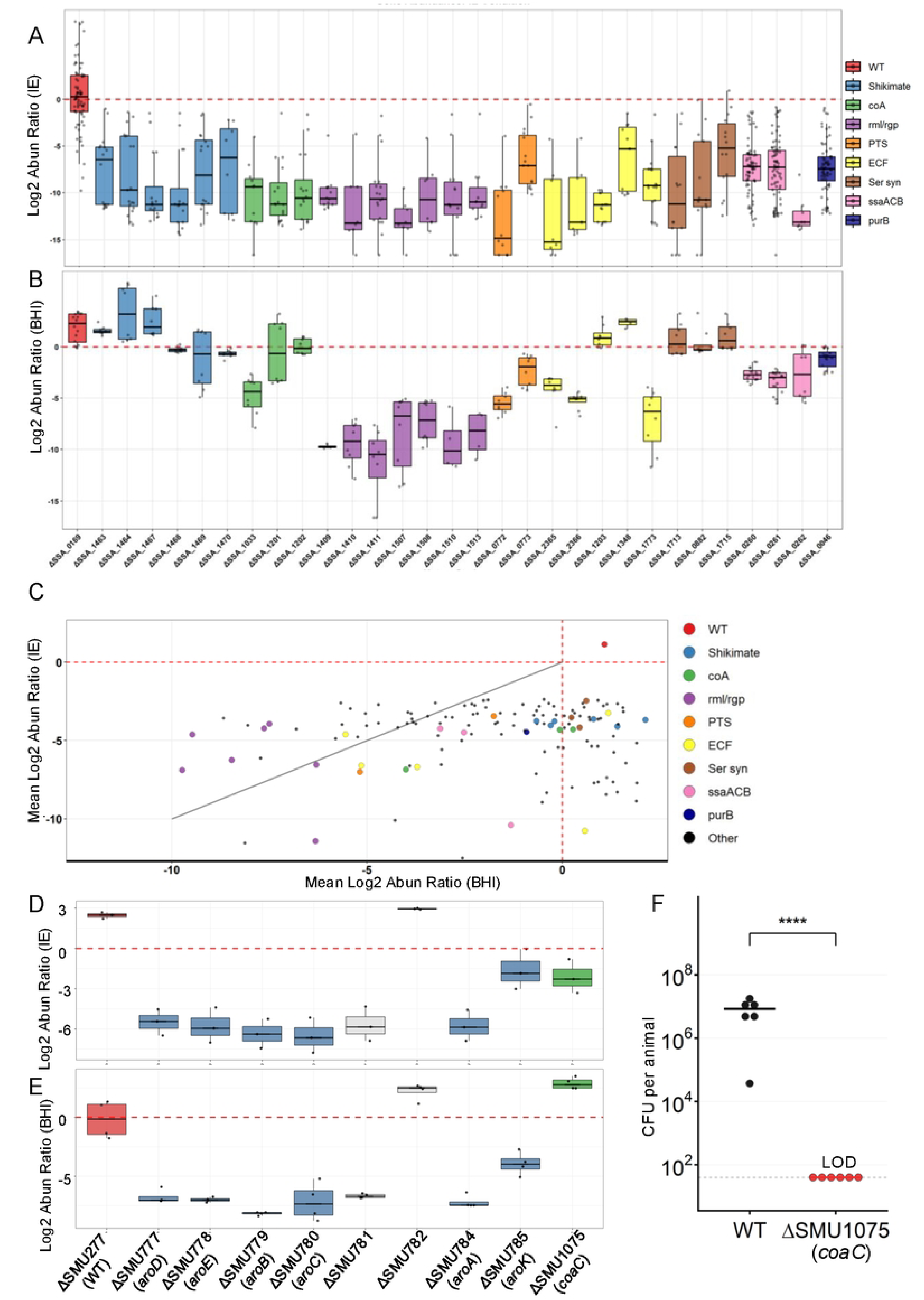
Fitness of selected mutants of S. *sanguinis* SK36 and S. *muttants* UAI 59 in the IE model and the rich medium BHI. **A.** Logi abundance ratios of selected S. *sanguinis* mutants in the output pool (IE model) compared to the input pool (inocula) across different animals. Each point represents a value of a mutant tested in one animal. For mutants with zero sequencing read counts, the abundance ratios were set to 0.0000 I for plotting log_2_ values. Dashed red line indicates an abundance ratio of 1. Abun, abundance. **B.** Log_2_ abundance ratios of selected S. *sanguinis* mutants in the output pool (BHI) relative to the input pool (inocula) across different BHI replicates. Each point represents a value of a mutant tested in one BHI replicate. Zero sequencing read counts were set to 0.00001 of abundance ratio for plotting. Dashed red line indicates an abundance ratio of 1. **C.** Pairwise comparison of mean Log_2_ abundance ratios of selected S. *sa11g11i11is* mutants in the IE model and BHJ. Each point represents the average Logi abundance ratio across multiple experiments. Dashed red lines indicate an abundance ratio of 1. The solid gray diagonal line represents equal fitness in both conditions; mutants plotted above the line exhibit a more pronounced fitness defect in BHl than in the IE model, while those below the line are more deficient in the IE model than in BHI. **D.** Log_2_ abundance ratios of S. *,nutans* mutants in the output pool (IE model) relative to the input pool across different animal replicates. Each point represents a single n1utant measured in one animal. Dashed red line indicates an abundance ratio of 1. **E.** Log_2_ abundance ratios of S. mu*tans* mutants in the output pool relative to the input pool across BHl replicates. Each point represents a single mutant measured in one BHI replicate. Dashed red line indicates an abundance ratio of 1. **F.** CFU per animal in vegetations collected 20 hours after inoculation with *ΔcoaC* and the wild type (WT) strain of *S. mutans* in the IE model. Note that the CFU for *ΔcoaC* was set at the limit of detection (LOD) since no colonies were recovered while assuming one colony formed without dilution. The horizontal bar represents the median of each strain, and the asterisk indicates statistical significance (P < 0.05) by Welch’s t-test after log transformation.

### *In vitro* growth assay of fitness factor mutants

To determine whether the observed IE fitness defects reflected general growth impairment or niche-specific requirements, we evaluated *in vitro* fitness in rich medium (BHI) using pooled ORFseq assays. To simulate the physiological environment of our vertebrate IE model, we utilized conditions representing the oxygen level of the left side of the heart (12% O_2_)[24, 37] and the elevated temperature observed during active infection (39°C) (Fig. S4). These analyses included three pools from the original IE fitness screen (final pool_1–3 Table S2) and a newly generated reconfirmation pool (Tables S8–S10). Of the 146 IE fitness determinant mutants identified *in vivo*, 137 were represented across the four pools and evaluated for growth (Figure 3B and 3C). Among these, 65 mutants exhibited reduced competitive fitness in BHI (abundance ratio <0.5) (Figure 3C), including two genes annotated as hypothetical proteins.

Growth phenotypes frequently diverged among genes within the same pathway. For example, *coaA* mutants exhibited severe growth defects in BHI, whereas downstream CoA biosynthetic mutants (Δ*coaB* and Δ*coaC*) did not (Figure 3B). Similar dissociations between *in vivo* fitness and *in vitro* growth were observed across multiple systems.

Genes encoding the shikimate pathway (*aroB, aroD, aroE, aroK, aroA,* and *aroC*) formed a contiguous cluster and were all required for IE fitness, yet none exhibited growth defects in BHI, indicating a niche-specific requirement for aromatic amino acid biosynthesis during IE (Figures 3A and 3B; Tables S8–S10). In contrast, rhamnan biosynthesis genes (*rlmABCD* and *rgpABCDF*) were required for both IE fitness and *in vitro* growth, consistent with a central role in cell wall integrity.

Core PTS components (EI and HPr) and select EII transporters also contributed to IE fitness, with some mutants displaying growth defects in BHI, suggesting roles in both general metabolism and host adaptation. Similarly, ECF transporters showed substrate-specific contributions to IE fitness, with only a subset required for *in vitro* growth. All three genes involved in serine biosynthesis (*serA, serB,* and *serC*) were required for IE fitness but dispensable for growth in BHI, consistent with sufficient serine availability in rich medium but not within cardiac vegetations.

Finally, mutants lacking components of the SsaACB Mn²⁺ transporter exhibited reduced fitness in BHI, differing from previous observations[36], likely reflecting the competition assay and the higher temperature and oxygen levels used in this study (Figure 3B; Tables S7–S9).

Together, these results demonstrate that IE fitness is supported by a combination of pathways required for general bacterial growth and others uniquely required in the host niche, underscoring the importance of metabolic capabilities.

### Assessment of the contribution of CoA and shikimate genes to growth and IE fitness in *S. mutans* UA159

To determine whether IE fitness determinants identified in *S. sanguinis* SK36 are conserved across IE-associated streptococcal species, we used the IE animal model to examine mutants of selected genes in *S. mutans*, an abundant oral species that is phylogenetically distant from *S. sanguinis* and exhibits antagonistic interactions with it in the oral environment [38–41]. We focused on genes involved in CoA synthesis and the shikimate pathway, as deletion of all but one, *coaA* (SSA_1033), produced no detectable growth defect in BHI in *S. sanguinis* SK36 (Figure 3B). Using targeted deletion, we successfully generated a *coaC* mutant and deletion mutants of all eight genes within the shikimate gene cluster, including six key shikimate pathway genes (*aroA, aroB, aroC, aroD, aroE,* and *aroK*), despite these genes having been previously reported as essential[42].

The generated mutants were pooled at comparable cell densities, together with the SMU277 deletion strain—which shows normal growth in BHI and was used as a wild-type (WT) reference, analogous to ΔSSA_0169 in *S. sanguinis* SK36. While ΔSMU277 was recovered in abundance from the *in vivo* IE model (Figure 3D; Table S11), the only other tested strain with high recovery was the mutant deleted for SMU782, encoding a YlbF/YmcA family competence regulator. The SMU782 result was consistent with that observed for the *S. sanguinis* strain deleted for the orthologous gene, SSA_1465. In contrast, SMU781, which encodes prephenate dehydrogenase, displayed reduced fitness in the IE model, differing from the outcome in *S. sanguinis* for the mutant of the orthologous gene, SSA_1466 (Tables S4 and S5). Pooled ORFseq assays performed in BHI medium *in vitro* using the same mutant pool revealed that the control strain, ΔSMU277, the Δ*coaC* mutant, and the ΔSMU782 mutant all displayed abundant growth in BHI, whereas the remaining six shikimate cluster mutants displayed reduced growth in BHI (Figure. 3E; Table S11).

To independently validate the ORFseq result in *S. mutans*, we performed a co-inoculation competition assay between the *coaC* mutant—which showed no growth defect in BHI—and the WT strain in the IE model. While the WT strain demonstrated robust recovery, no colonies of the *coaC* mutant were recovered, confirming a severe IE-specific fitness defect (Figure 3F), consistent with the pooled ORFseq findings.

### The growth defect of shikimate pathway mutants can be restored by increased peptide transport in *S. mutans*

To more deeply investigate additional IE fitness determinants, we next focused on the shikimate pathway. To determine how disruption of the shikimate pathway impacts IE fitness in *S. mutans*, we screened for compensatory mutations that could restore growth in shikimate pathway mutants, which exhibited strong growth defects in BHI medium for seven of the eight genes tested. Serial passage experiments were performed using newly generated *S. mutans* UA159 shikimate gene deletion mutants, with ΔSMU782—which showed no growth defect in BHI—used as a control. For most genes, two independently evolved populations were sequenced, while four populations were sequenced for ΔSMU785. Whole-genome sequencing revealed that the evolved mutant populations of four genes whose deletion resulted in growth defects acquired duplications encompassing a peptide transporter gene cluster (SMU255 to SMU259) (Figures 4A to 4D), suggesting that increased peptide uptake can compensate for loss of shikimate pathway function in *S. mutans*.

**Figure 4.**
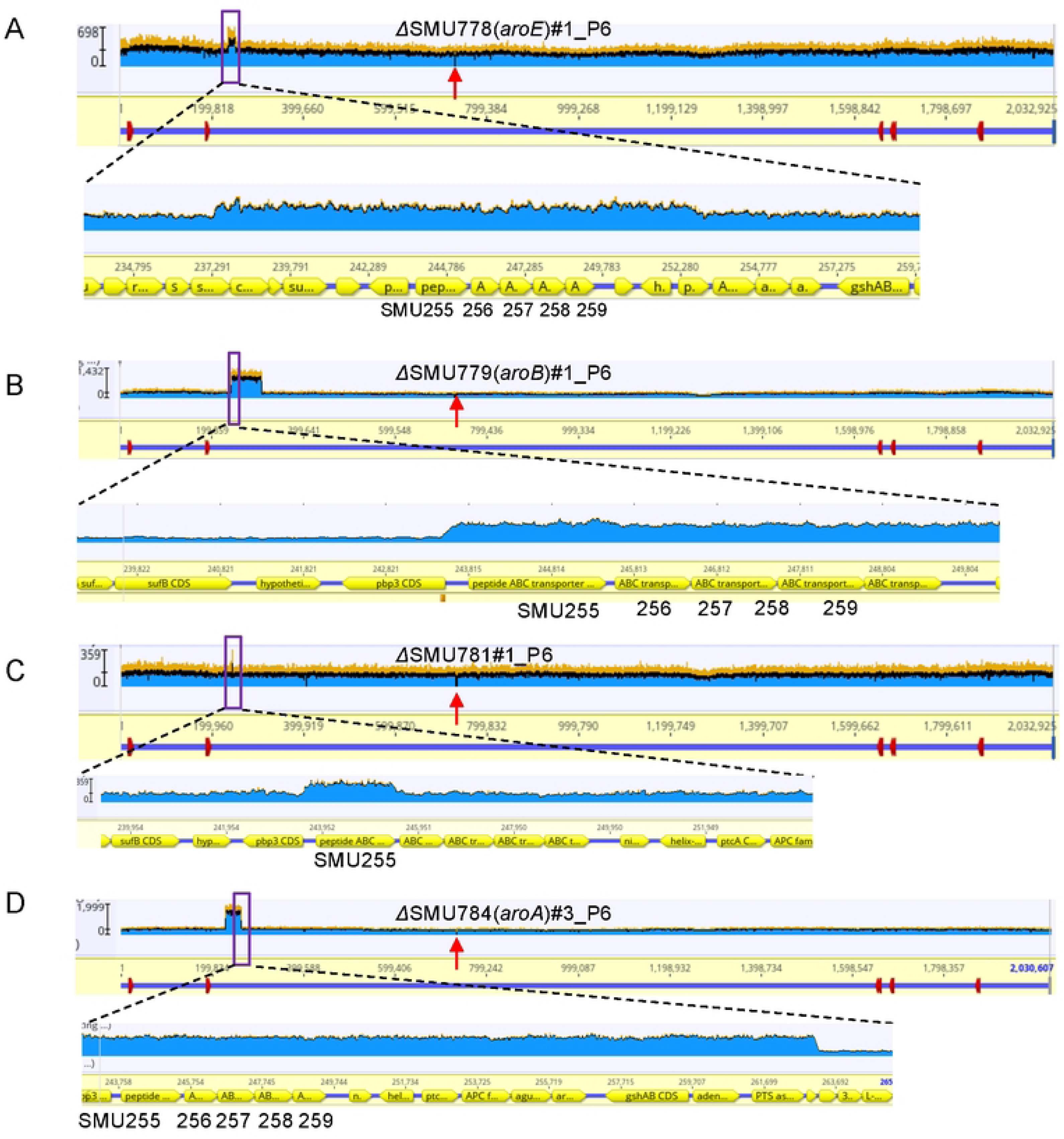
Compensatory mutations of 1E fitness factor mutants in the S. *mutants* shikimate gene cluster. **A-D.** Gene duplications arising within the peptide ABC transporter gene cluster from four independently evolved IE fitness factor mutants in the shikimate gene cluster (SMU778, SMU779, SMU781 and SMU784) of the populations SMU778#1 **(A),** SMU779#1 (B), SMU781#1 **(C)** and SMU784#3 **(D).** The purple boxes indicate the enlarged peptide ABC transporter gene cluster. Vertical red arrow, site of original gene deletion; the height of the blue segn1ents indicates the number of sequence reads mapped to the reference sequence at the coordinates shown on the X axis. A doubling of sequence reads indicates a duplication of the affected region.

### Competitiveness of *S. sanguinis* shikimate pathway cluster mutants in BHI and rabbit serum

To further validate the results in the ORFseq in BHI (Figure 3B) and to evaluate mutant behavior in rabbit serum, competition experiments were performed for eight *S. sanguinis* mutants within the shikimate gene cluster. These included six shikimate pathway genes identified as IE fitness factors. The WT-like strain ΔSSA_0169 was used as a control, and all assays were performed under physiological oxygen levels representative of the left side of the heart (12% O₂) and at 39°C. For competition assays, each mutant strain, as well as the control ΔSSA_0169 strain carrying a kanamycin resistance cassette, was mixed at approximately equal cell numbers with the reference strain JFP36, which harbors an erythromycin resistance cassette inserted at the SSA_0169 locus, and CFUs were enumerated after 24 hours of co-culture. In BHI medium and rabbit serum, mutants deleted for each of the seven genes from SSA_1463 through SSA_1469 showed no detectable fitness defect under either condition (Fig. S5A and S5B). In contrast, ΔSSA_1470 exhibited reduced fitness in rabbit serum but not in BHI medium (Fig. S5A and S5B).

### The *in vitro* growth defect of *S. sanguinis* CoA mutants can be restored by increased fatty acid synthesis

To investigate the mechanism by which genes contributed to IE fitness, we first focused on CoA biosynthesis in *S. sanguinis*. Three CoA synthesis genes—*coaA* (SSA_1033), *coaB* (SSA_1201), and *coaC* (SSA_1202)—were examined using targeted deletion mutants to assess their fitness in BHI medium and rabbit serum; the WT-like strain ΔSSA_0169 was used as a control, and all experiments followed the same design used for shikimate pathway mutants above. Consistent with pooled ORFseq results (Figure 3B), the Δ*coaA* mutant exhibited reduced growth in BHI and Δ*coaB* and Δ*coaC* showed no growth defect (Figure 5A). In addition, in rabbit serum, Δ*coaA*, Δ*coaB* and Δ*coaC* displayed reduced fitness (Figure 5B).

**Figure 5.**
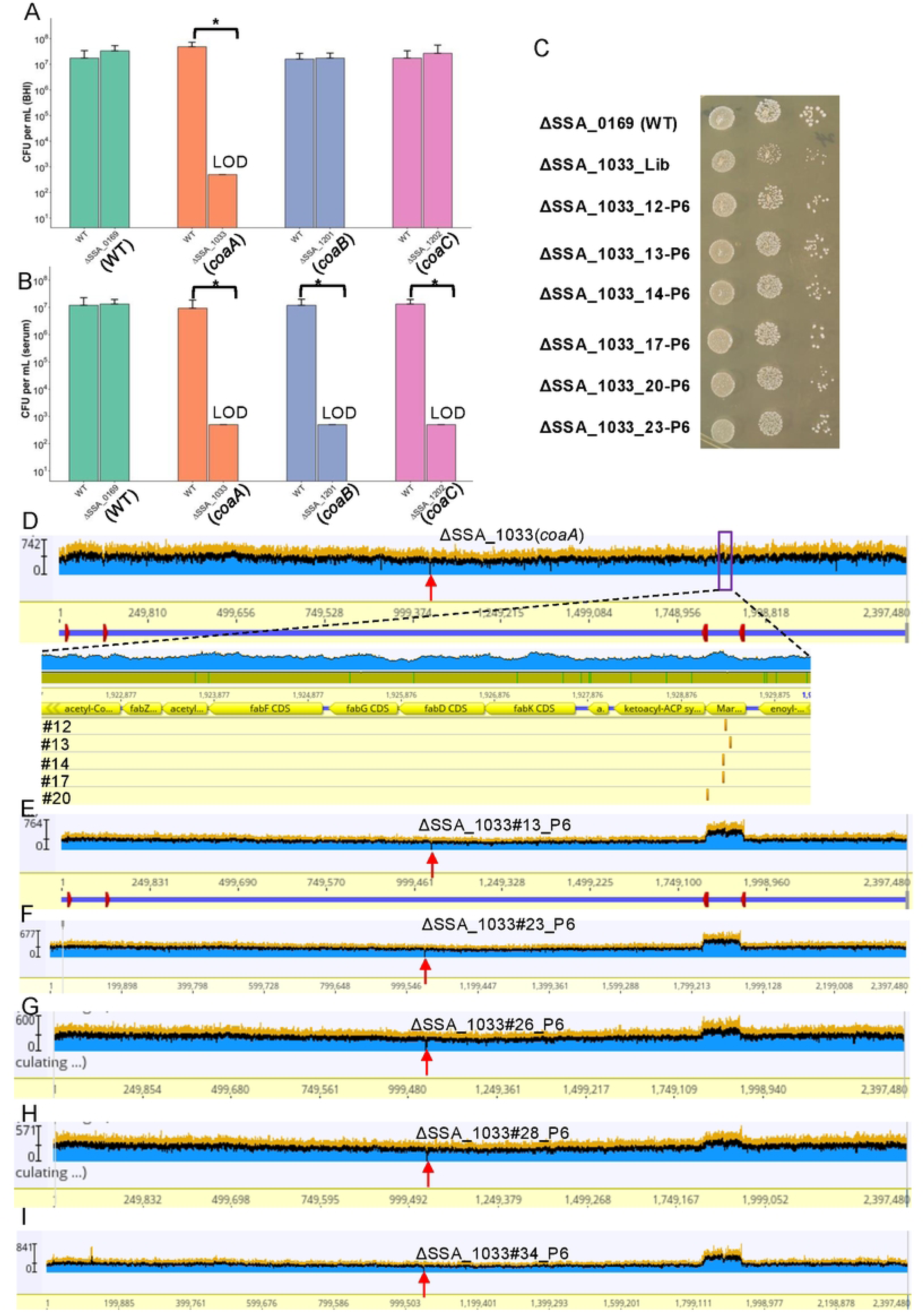
Growth phenotypes and compensatory mutations of CoA-biosynthesis IE fitness factor mutants in S. *sanguinis.* **A-B.** Competitive growth of selected mutants relative to the JFP36 strain in BHI medium (A) and rabbit serun1 (B). Bars represent CFU 1neasured 24 hours after coculture of 111utants with the JFP36 wild-type strain. The li111it of detection (LOO) indicates the minimu111 detectable CFU, assuming one colony formed without dilution since no colonies were recovered. The asterisk indicates statistical significance (P < 0.05) by Welch’s t-test after log transformation. **C.** Growth of WT, the *ΔcoaA* mutant library stock and the evolved *ΔcoaA* mutants. Growth of the strains indicated. Two microliters of cells at an 0D_600_ of I was diluted 20-fold (column !), or 400-fold (column 2), or 8,000-fold (column 3) and then spotted onto BHI-agar and allowed to grow anaerobically for 3 days. **D.** Suppressor n1utations arising in the FabT transcriptional repressor from five independently evolved *ΔcoaA* mutants (#12, #13, # 14, #17 and #20). Purple boxes indicate the enlarged fatty acid synthesis (F ASII) gene cluster. The rRNA operons are indicated by red boxes in panel. The height of the blue segments indicates the number of sequence reads mapped to the reference sequence at the coordinates shown on the X axis. The vertical arrow in red indicates the deleted *coaA* gene. The vertical gold bars indicate the positions of the n1utations in *thefabT* gene. **E-I.** Gene duplications arising within the F ASU gene cluster regulated by FabT from three independently evolved *ΔcoaA* mutant populations #13 **(E),** #23 **(F),** #26 **(G), #28 (H)** and #34 **(I).** Note that the duplicated regions are flanked by directly repeated rRNA operons (indicated by red boxes in panel E). Vertical red an-ow, site of original gene deletion; the height of the blue segments indicates the number of sequence reads mapped to the reference sequence at the coordinates shown on the X axis. A doubling of sequence reads indicates a duplication of the affected region.

To explore how altered CoA synthesis might impact IE survival, we screened for compensatory mutations that could restore growth in the *S. sanguinis* Δ*coaA* mutant, which displayed a strong growth defect in BHI medium. Newly generated Δ*coaA* mutant populations were subjected to serial passage for six cycles, resulting in multiple independently evolved populations. Comparative growth analysis showed that the evolved Δ*coaA* populations formed larger colonies and exhibited markedly improved growth in BHI medium relative to the unevolved library strain (Figure 5C). Whole-genome sequencing revealed that five populations (#12, #13, #14, #17, and #20) acquired mutations in *fabT* (Figure 5D), a negative transcriptional regulator of the type II fatty acid synthesis (FASII) pathway[43], including one truncation mutation in population #20. In addition, five populations (#13, #23, #26, #28, and #34) exhibited duplications of the FASII gene cluster, with population #13 harboring both a *fabT* mutation and FASII duplication (Figures 5D to 5I). Sequencing of the original Δ*coaA* mutant from the ORFseq library confirmed the absence of secondary mutations. Together, these results suggest that increased FASII expression can compensate for impaired CoA synthesis, linking CoA availability to fatty acid biosynthesis.

## Discussion

In this study, we performed the first genome-wide *in vivo* analysis of bacterial fitness in IE, systematically defining the genetic requirements for *S. sanguinis* growth and proliferation in a vertebrate IE model (Figure 1). By combining a comprehensive mutant library with optimized pooled screening and rigorous *in vivo* validation, we identified 146 genes required for IE fitness (Table S6), the vast majority of which had not previously been associated with endocarditis in any bacterium. Although this systems-level approach provides a broad landscape of infection, we acknowledge the inherent limitations of pooled screenings. In addition, genetic redundancy can mask the importance of specific pathways even in single-strain inoculations. Thus, while our screen emphasizes metabolic and fitness networks, it does not preclude the existence of traditional virulence factors that may fall outside the detection limits of this methodology.

To assess the reliability of our screening approach, we first compared the 146 identified fitness factors with those previously reported by our group. Earlier studies demonstrated reduced fitness following deletion of individual open reading frames (ORFs) such as *purB*[19, 44], *thrB*[19, 44], *bacA*[19], *ssaB*[20, 21], *sodA*[21], *nox*[22], and *trxR1*[23] and upon simultaneous deletion of multiple ORFs, including Δ*ssaACB*[24, 25], Δ*ecf* (ΔSSA_2365-SSA_2367)[31], and Δ*nrdHEKF*[23]. The *nrdHEKF* and *trxR1*[23] genes are essential under oxic conditions and thus, were not included in this study. Our results demonstrate strong concordance with prior findings. The screen confirmed nine previously identified IE fitness determinants, including SSA_1044 (*thrB*)[19, 44], SSA_1127 (*nox*)[22], SSA_1959 (*bacA*)[19], and SSA_0260 (*ssaB*)[20, 21], as well as two components of the ECF transporter system: T (SSA_2365) and A1 (SSA_2366) from Δ*ecf* (ΔSSA_2365-SSA_2367). Therefore, all previously reported determinants were validated except *sodA*, which showed significantly reduced fitness in three of four experiments but an abundance ratio near 1 in one experiment. Our earlier work showed that *sodA* mutation impaired IE fitness, although less severely than *ssaB* mutation [21], which could explain our current results. Furthermore, consistent with a prior signature-tagged mutagenesis study that detected no significant fitness defects among mutants targeting 33 predicted cell-wall–anchored proteins or their sortases[28], the corresponding ORFseq mutants in the present screen did not exhibit reduced fitness (Tables S5 and S6). This agreement across distinct methodologies reinforces the robustness of our findings.

We also compared our findings with those reported previously by other groups who worked with *S. sanguinis*. The first of these is *nt5e* (SSA_1234), which encodes a surface protein capable of hydrolyzing extracellular ATP[45]. Because this activity likely functions as a “public good,” mutants deficient in *nt5e* would not be expected to show reduced fitness in pooled screens[46]. Consistent with this idea, reduced fitness of the *nt5e* mutant was demonstrated when mutant, wild-type, and complemented strains were inoculated into separate animals[45]. Other studies identified *mur2* (presumably SSA_1095), SSA_1099[26], and the type IV pilus gene *pilF* (SSA_2318)[27] as IE fitness factors. Discrepancies between these studies and ours may reflect differences in pooled versus individual mutant testing, variation in rabbit IE models, or differences in strain backgrounds. Notably, reduced fitness of the *pilF* mutant was demonstrated relative to SK36 variants selected for twitching motility[27], leaving unclear whether the mutant would have shown reduced fitness compared to wild-type SK36.

The identified IE fitness determinants span core processes (Figure 3A and Fig. S3) including central metabolism, cell envelope biogenesis, transport, and information processing, revealing that IE fitness depends on the coordinated function of multiple interconnected pathways. Notably, several pathways—such as coenzyme A biosynthesis, rhamnan-mediated cell wall assembly, and the shikimate pathway—were represented almost in their entirety, underscoring the depth and completeness of the screen.

Comparative analysis demonstrated that most IE fitness genes are present in major IE pathogens, including *Enterococcus*, *Staphylococcus*, and other *Streptococcus* species. Demonstration that a subset of these genes is also required for IE in the distantly related oral species *S. mutans* suggests that these selected determinants are important for IE fitness in other oral streptococcal species as well (Table S6). Importantly, several fitness determinants, exemplified by *coaC*, were required specifically *in vivo* or in serum but not under standard laboratory growth conditions, highlighting the limitations of *in vitro* screens for identifying clinically relevant fitness factors. Other pathways exhibited species-specific dependencies, underscoring the context dependence of bacterial fitness strategies.

Beyond defining infection-associated fitness requirements, our findings identify multiple pathways that represent promising targets for antimicrobial development. Several IE-associated fitness systems—including the shikimate pathway, CoA biosynthesis, rhamnan cell wall assembly, PTS components, ECF transporters, serine biosynthesis, and manganese acquisition through SsaACB—are absent or highly divergent in mammalian hosts, enhancing their therapeutic selectivity. The identification of multiple genes within individual pathways or protein complexes (Figure 3b and Fig. S3) as IE fitness determinants suggests that these systems function as coordinated biological modules, which may increase their susceptibility to pharmacological inhibition. Among these, the shikimate pathway represents an especially attractive target for IE control. This pathway is completely absent in humans, minimizing the potential for host toxicity[47, 48], yet is highly conserved across diverse bacterial pathogens, including major etiologic agents of infective endocarditis (Table S6). Moreover, the enzymatic reactions and structural features of shikimate pathway proteins have been extensively characterized[49], providing well-defined and drug-targetable enzymatic steps that facilitate rational antimicrobial design[47, 48]. Several inhibitors targeting this pathway already exist, for example, glyphosate inhibits 5-enolpyruvylshikimate-3-phosphate synthase, and multiple shikimate analogs have demonstrated antimicrobial or antiparasitic activity, including compounds developed against malaria parasites[50, 51]. Similarly, enzymes involved in CoA biosynthesis[52] and ECF transporter systems[53, 54] represent promising antimicrobial targets due to their central roles in metabolism and micronutrient acquisition. Rhamnan biosynthesis enzymes offer potential targets for disrupting streptococcal cell wall integrity[55], while manganese transport systems such as SsaACB play critical roles in oxidative stress resistance and host adaptation[21, 36]. These findings collectively highlight infection-specific metabolic and transport pathways as vulnerable nodes in bacterial pathogenesis and provide a rational framework for developing targeted strategies to prevent or treat infective endocarditis.

By integrating experimental evolution, we demonstrate that IE fitness networks are highly adaptable. For instance, the disruption of key pathways—such as coenzyme A biosynthesis in *S. sanguinis*—triggered reproducible compensatory adaptations through the upregulation of fatty acid synthesis, revealing extensive metabolic plasticity. Similarly, experimental evolution of *S. mutans* shikimate mutants revealed recurrent duplication of peptide transporter genes supporting fitness under these conditions. Collectively, these findings expand current understanding of the genetic and functional basis of streptococcal fitness during infective endocarditis and provides a resource for future studies investigating bacterial adaptation and host-associated growth requirements. Continued investigation of these fitness determinants may help identify pathways that influence bacterial persistence during infection and could inform future preventive or therapeutic strategies.

## Materials and methods

### Strains

The 2,048 non-essential mutant library used in this study was sourced from a previous study[30]. Additionally, 27 knockout mutants of newly annotated open reading frames were generated using the same method as in the previous study[30], while Δf1fo mutants were generated in our recent study[32] (Table S1).

### Mutant pooling

All mutants were individually preserved in 20% glycerol stocks and stored at -80°C until use. Selected mutants were inoculated from the -80°C glycerol stock into Eppendorf tubes containing 300 μL of BHI medium and grown overnight under microaerobic conditions (6% O₂, 7.2% CO₂, 7.2% H₂, and 79.6% N₂) using an Anoxomat (Advanced Instruments, Norwood, MA) jar at 37°C. Following overnight growth, the 300 μL cultures were transferred into 1.2 mL of fresh BHI medium and cultured for an additional 3 hours under the same microaerobic conditions. A 100 μL sample was collected to measure cell density by optical density at 600 nm (OD_600_). Equal OD_600_ values of different mutants were pooled to create input pools (Figure 1).

The cell cultures were then pelleted by centrifugation at 3,000 rpm for 10 minutes at room temperature, and the supernatant was discarded. The resulting 3 mL pellet was thoroughly mixed with 1 mL of 80% glycerol, aliquoted, and stored at -80°C until further use. For preparing inoculum, the 1 mL glycerol stock of a mixed mutant pool was washed twice in 10 mL PBS and adjusted to OD_600_ of 0.8, approximating 10^8^ CFU per mL.

### IE animal model

To assess fitness, an endocarditis model was employed as described previously[24] with modifications. In brief, New Zealand White rabbits were sedated, anesthetized, and provided an extended-release analgesic prior to the procedure. Endocardial damage was induced by inserting a PE-90 catheter into the right carotid artery until it met or passed a short distance through the aortic valve, with placement monitored via ultrasound imaging. The catheter was sealed and the incision site was then sutured closed. After a two-day recovery period, sedated rabbits were inoculated with 0.5 ml of a pooled mutant strain suspension prepared as described above via a peripheral ear vein. Approximately 20 hours post-inoculation, the rabbits were sedated and euthanized through intravenous administration of Euthasol. Cardiac vegetations were collected and homogenized in PBS.

### DNA isolation

Genomic DNA (gDNA) was isolated from the homogenized vegetation samples and from the inocula. Briefly, cells were pelleted by centrifugation at 10,000 rpm for 10 minutes at room temperature, then resuspended in 200 μL of resuspension buffer (20 mM EDTA, 200 mM Tris-HCl, 2% Triton X-100). For lysis, 200 μL of AL lysis buffer (Qiagen, 19075) was added, and the mixture was incubated for 1 hour at room temperature. DNA was precipitated by adding 1 mL of 100% ethanol containing 100 mM sodium acetate. Following washing and drying, the DNA was resuspended in 150 μL of water and prepared for ORFseq library construction.

### ORFseq library preparation

ORFseq library preparation followed a modification of a previously described protocol[31]. Genomic DNA (gDNA) was fragmented to approximately 500 bp using a Covaris S2 Ultrasonicator under the following settings: Duty cycle - 5%; Intensity - 5.0; Bursts per second - 200; Power - 23 W; Mode - Frequency sweeping; Treatment time - 1/2 = 60 sec/40 sec. PolyC tails were then added to the 3ʹ ends of the fragmented DNA using terminal deoxynucleotidyl transferase (Promega, USA) at 37°C for one hour, followed by enzyme inactivation at 75°C for 20 minutes.

PolyC-tailed DNA fragments were purified with AMPure XP beads (Beckman, USA) and used as templates in PCR amplification with Platinum™ Taq DNA Polymerase (Invitrogen, 10966026).The first round of PCR was performed with primers olj376 and K10_Truseq (Table S12) under the following conditions: an initial denaturation at 94°C for 2 minutes, followed by 25 cycles of 94°C for 30 seconds, 60°C for 30 seconds, and 68°C for 30 seconds, with a final extension at 68°C for 5 minutes, and a hold at 4°C. PCR products were then purified using AMPure XP beads. A second PCR round was performed using PE1npKan as the universal 3ʹ primer (Table S12) and distinct Truseq_HT primers (Table S12) to index the samples. The PCR conditions were identical to the first round. The final PCR products were purified again with AMPure XP beads and submitted to the VCU DNA Core Facility for NGS sequencing.

### ORFseq library sequencing and quantification of mutants

Sequencing was conducted on the Illumina platform with 100 cycles of single-end sequencing. The leading sequence (TTTTAGTACCTGGAGGGAATAATG), corresponding to the 3’-end of the *APH(3’)-IIIa* (kanamycin resistance) gene, was trimmed from the reads using Cutadapt[56]. For the ORFseq for *S. sanguinis*, the trimmed reads were then aligned to the reference *S. sanguinis* SK36 genome (CP071435.1) using Bowtie2[57], and read counts were calculated with featureCounts[58]. For ORFseq in *S. mutans*, the trimmed reads were aligned to the reference *S. mutans* UA159 genome (NC_004350.2) using Bowtie2, and read counts were calculated based on base coverage within the corresponding genes. Mutant abundance was calculated by averaging two to three technical replicates from input or output samples following normalization by total read counts. The abundance ratio for each biological replicate was calculated by dividing output abundance by input abundance. The overall abundance ratio was determined by averaging abundance ratios across biological replicates.

### Screening workflow

The fitness factor screenings were conducted in three stages (Stages I–III). In Stage I, seventeen initial mutant pools (“initial screen pool-1” to “initial screen pool-17”) were tested. Mutants showing significant fitness differences in Stage I were retested in Stage II using three re-screen pools (“re-screen pool-1” to “re-screen pool-3”) (Figure 1B). In Stage III, candidates with abundance ratios significantly different from 1 were further validated and compared directly using four final pools (“final pool-1” to “final pool-4”), enabling relative fitness comparisons among candidates. This stage involved testing in four final pools (“final pool-1” to “final pool-4”) in which the relative fitness of each candidate could be compared with the others. In total, the screening process involved 24 mixed mutant pools, yielding 3,435 tests for 2,039 unique ORF deletion mutants. Of these, 1,210 mutants were tested once with the majority identified as fitness normal, 829 mutants (40.6%) were tested in at least two experiments, while 37 mutants remained untested.

### Selection criteria for fitness factor candidates

Fitness factor candidates were identified based on *p*-values, abundance ratios, and number of tests performed per mutant, as follows.1) Single-Experiment Test Mutants: Mutants tested once were considered fitness-reduced if the abundance ratio was significantly different from 1 and the ratio was < 0.025. This value was chosen because it was half the average ratio for the three fitness-reduced control mutants (ΔSSA_0046, ΔSSA_0260 and ΔSSA_0261). 2) Two-Experiment Test Mutants: Mutants tested twice were classified as fitness-reduced if both results were significant and the overall abundance ratio was < 0.2. This was the rounded highest value for the fitness-reduced controls. 3) Three or More-Experiment Test mutants: Mutants tested three or more times were considered fitness-reduced if the number of significant tests exceeded non-significant ones, all significant tests showed reduced abundance, and the overall ratio was < 0.2.

### *In vitro* competition assay

JFP36 (Erm^r^; ΔSSA_0169::p*Serm*; 19423626) and selected Km-resistant mutants were cultured separately from -80°C glycerol stocks in 2 mL of BHI medium without antibiotics in 4-mL tubes. The cultures were incubated overnight (∼16 hours) at 37°C under microaerobic conditions (6% O₂). The next day, cultures of these strains were diluted 1:1,000,000 in BHI or serum. Each of the Km-resistant mutants was then mixed with JFP36 at a 1:1 ratio. Colony-forming units (CFUs) of the 1:1 mixtures were determined at time 0 by plating on BHI-agar containing either 10 μg/mL Erm or 500 μg/mL Km. The mixtures were then incubated for 24 hours at 39°C under 12% O₂. After 24 hours of growth, CFUs were measured again by plating on BHI-agar containing 10 μg/mL Erm or 500 μg/mL Km.

### Identification of compensatory mutations

Antibiotic-resistant colonies selected on agar plates were inoculated into 1 mL of BHI broth containing the appropriate antibiotics and incubated anaerobically at 37°C for 2-3 days. Cultures were grown until the OD_600_ reached 0.1–0.5; this initial culture was designated P0. For the first passage (P1), 300 µL of the P0 culture was transferred into 3 mL of BHI containing antibiotics and grown to saturation. The P1 culture was mixed thoroughly by pipetting five times with a P1000 pipette, and 50 µL was then transferred into 1 mL of fresh BHI with antibiotics to initiate P2. Cultures were grown to an OD_600_ of 0.1–0.5 before the next transfer. This sequential passaging procedure was repeated through P6. Passages P2–P5 were each grown in 1 mL volumes, whereas the final passage (P6) was grown in 3 mL. Aliquots of P1 and P6 cultures were preserved at −80°C in BHI supplemented with 20% glycerol (final concentration). DNA extraction and whole-genome sequencing were performed using cells from P6. For variant calling, whole-genome sequencing was carried out by SeqCenter (https://www.seqcenter.com/) using the shotgun method with 2 × 150 paired-end sequencing.

Fastq files were aligned to an updated SK36 reference genome sequence (CP071435.1) or *S. mutans* UA159 genome (NC_004350.2) using Geneious Prime software (https://www.geneious.com/) after trimming via the BBDuk method. Sequences with an average coverage of ≥100 were used for subsequent analysis. Variations in the genome were exported from Geneious Prime. To identify the mutated segment, the frequencies (percentages) of all mutations belonging to a certain segment were determined using the procedure recommended by the makers of Geneious Prime.

### Statistics

Statistical analysis. Statistical analyses were performed using Microsoft Excel. Unless otherwise indicated, comparisons of competitiveness between two strains were conducted using two-tailed Welch’s t-tests. Statistical details of specific tests used, are provided in the corresponding figure legends. A P value of <0.05 was considered statistically significant.

## Data availability

All data needed to evaluate the conclusions in the paper are present in the paper and/or the supplemental files. The genome sequence data for the suppressor identification were deposited to GenBank (PRJNA1431959). All the transgenic materials, including the essential gene deletion mutants, are available from the authors upon request.

## Contributions

P.X. conceived the project, and P.X., L.B. and T.K. designed the experiments. L.B., J.B. and J.A.V. performed animal surgeries and collected vegetations with the help of V.A., V.F.A., J.S.B and N.Z.. L.B., K.M.T., P.X. and T.K. analyzed data. L.B., V.F.A., V.A., Z.Z. and N.Z. prepared bacterial pools. L.B. and V.A. generated new mutants. L.B., T.K. and N.Z. performed *in vitro* studies. L.B. prepared bacterial inocula, extracted gDNA, constructed ORFseq libraries, and wrote the original manuscript. L.B., P.X. and T.K. revised the manuscript and all authors approved it.

## Acknowledgements

We would like to thank Dr. Yan Bao at the School of Agriculture and Biology, Shanghai Jiao Tong University, for discussions regarding experimental design and critical reading of the manuscript. We also thank Dr. Junling Ren and Dr. Huizhi Wang at the Philips Institute for Oral Health Research, Virginia Commonwealth University, for helpful discussion during the project. We are grateful to Dr. Seon-Sook An of the Philips Institute for Oral Health Research, Virginia Commonwealth University for assistance with animal experiments, Vladimir Lee, Linyin Xie and Kalyan Maliempati for genomic sequencing services, Katherine Atran, Danielle Dexter Keeton, Magen Lindsey, and Kali Williams at the Division of Animal Resources, Virginia Commonwealth University for assistance with animal surgery, and Dr. Krista Scoggins, and Mahesh Jonnalagadda, for animal care and welfare. Services in support of the presented research were partially provided by the VCU Massey Comprehensive Cancer Center Bioinformatics Shared Resource. Massey is supported, in part, with funding from NIH-NCI Cancer Center Support Grant P30 CA016059. This work was funded by grants from the National Institutes of Health R01DE030121 (P.X. and T.K.), R03DE034511 (L.B.) and the CCTR Endowment Fund (P.X.). During the preparation of this work, the authors used ChatGPT-4 in order to improve readability and language. After using it, the authors reviewed and edited the content as needed and took full responsibility for the content of the publication.

## References

1. Holland, T.L., Baddour, L.M., Bayer, A.S., Hoen, B., Miro, J.M., and Fowler, V.G. (2016). Infective endocarditis. Nat Rev Dis Primers 2, 16059.

2. Iung, B., and Duval, X. (2019). Infective endocarditis: innovations in the management of an old disease. Nat Rev Cardiol 16, 623–635.

3. Li, M., Kim, J.B., Sastry, B.K.S., and Chen, M. (2024). Infective endocarditis. Lancet 404, 377–392.

4. Taduru, S.S. (2023). 30-Year Trends of Incidence and Mortality of Infective Endocarditis in the United States-Unveiling the Age- and Gender-Related and Regional Disparities. Am J Cardiol 204, 421–422.

5. Werdan, K., Dietz, S., Löffler, B., Niemann, S., Bushnaq, H., Silber, R.E., Peters, G., and Müller-Werdan, U. (2014). Mechanisms of infective endocarditis: pathogen-host interaction and risk states. Nat Rev Cardiol 11, 35–50.

6. McCormick, J.K., Tripp, T.J., Dunny, G.M., and Schlievert, P.M. (2002). Formation of vegetations during infective endocarditis excludes binding of bacterial-specific host antibodies to *Enterococcus faecalis*. J. Infect. Dis. 185, 994–997.

7. Durack, D.T. (1975). Experimental bacterial endocarditis. IV. Structure and evolution of very early lesions. J Pathol 115, 81–89.

8. Moreillon, P., and Que, Y.A. (2004). Infective endocarditis. Lancet 363, 139–149.

9. Strom, B.L., Abrutyn, E., Berlin, J.A., Kinman, J.L., Feldman, R.S., Stolley, P.D., Levison, M.E., Korzeniowski, O.M., and Kaye, D. (1998). Dental and cardiac risk factors for infective endocarditis. A population-based, case-control study. Ann Intern Med 129, 761–769.

10. Nobbs, A., and Kreth, J. (2019). Genetics of *sanguinis*-Group Streptococci in Health and Disease. Microbiol Spectr 7.

11. Havers-Borgersen, E., Østergaard, L., Holgersson, C.K., Stahl, A., Schmidt, M.R., Smerup, M., Køber, L., and Fosbøl, E.L. (2024). Infective endocarditis with or without congenital heart disease: clinical features and outcomes. Eur Heart J 45, 4704–4715.

12. Thornhill, M.H., Gibson, T.B., Yoon, F., Dayer, M.J., Prendergast, B.D., Lockhart, P.B., O’Gara, P.T., and Baddour, L.M. (2024). Endocarditis, invasive dental procedures, and antibiotic prophylaxis efficacy in US Medicaid patients. Oral Dis 30, 1591–1605.

13. Thornhill, M., Prendergast, B., Dayer, M., Frisby, A., Lockhart, P., and Baddour, L.M. (2024). New evidence calls into question NICE’s endocarditis prevention guidance. Br Dent J 236, 702–708.

14. Wilson, W.R., Gewitz, M., Lockhart, P.B., Bolger, A.F., DeSimone, D.C., Kazi, D.S., Couper, D.J., Beaton, A., Kilmartin, C., Miro, J.M., et al. (2021). Prevention of Viridans Group Streptococcal Infective Endocarditis: A Scientific Statement From the American Heart Association. Circulation 143, e963–e978.

15. Epprecht, J., Ledergerber, B., Frank, M., Greutmann, M., van Hemelrijck, M., Ilcheva, L., Padrutt, M., Stadlinger, B., Özcan, M., Carrel, T., et al. (2024). Increase in Oral Streptococcal Endocarditis Among Moderate-Risk Patients: Impact of Guideline Changes on Endocarditis Prevention. JACC Adv 3, 101266.

16. Bae, J., Park, J.H., Lee, M., Jo, H.J., Lee, C.M., Kang, C.K., Choe, P.G., Park, W.B., Kim, N.J., Kim, I., et al. (2024). Clinicomicrobiological risk factors for infective endocarditis in viridans group streptococci bacteraemia. J Antimicrob Chemother 79, 2327–2333.

17. Strange, J.E., Østergaard, L., Køber, L., Bundgaard, H., Iversen, K., Voldstedlund, M., Gislason, G.H., Olesen, J.B., and Fosbøl, E.L. (2023). Patient Characteristics, Microbiology, and Mortality of Infective Endocarditis After Transcatheter Aortic Valve Implantation. Clin. Infect. Dis. 77, 1617–1625.

18. Salgado-Pabón, W., and Schlievert, P.M. (2016). Aortic Valve Damage for the Study of Left-Sided, Native Valve Infective Endocarditis in Rabbits. Methods Mol Biol 1396, 73–80.

19. Paik, S., Senty, L., Das, S., Noe, J.C., Munro, C.L., and Kitten, T. (2005). Identification of virulence determinants for endocarditis in Streptococcus sanguinis by signature-tagged mutagenesis. Infect Immun 73, 6064–6074.

20. Das, S., Kanamoto, T., Ge, X., Xu, P., Unoki, T., Munro, C.L., and Kitten, T. (2009). Contribution of lipoproteins and lipoprotein processing to endocarditis virulence in Streptococcus sanguinis. J Bacteriol 191, 4166–4179.

21. Crump, K.E., Bainbridge, B., Brusko, S., Turner, L.S., Ge, X., Stone, V., Xu, P., and Kitten, T. (2014). The relationship of the lipoprotein SsaB, manganese and superoxide dismutase in Streptococcus sanguinis virulence for endocarditis. Mol Microbiol 92, 1243–1259.

22. Ge, X., Yu, Y., Zhang, M., Chen, L., Chen, W., Elrami, F., Kong, F., Kitten, T., and Xu, P. (2016). Involvement of NADH Oxidase in Competition and Endocarditis Virulence in Streptococcus sanguinis. Infect Immun 84, 1470–1477.

23. Rhodes, D.V., Crump, K.E., Makhlynets, O., Snyder, M., Ge, X., Xu, P., Stubbe, J., and Kitten, T. (2014). Genetic characterization and role in virulence of the ribonucleotide reductases of *Streptococcus sanguinis*. The Journal of biological chemistry 289, 6273–6287.

24. Baker, S.P., Nulton, T.J., and Kitten, T. (2019). Genomic, Phenotypic, and Virulence Analysis of Streptococcus sanguinis Oral and Infective-Endocarditis Isolates. Infect Immun 87.

25. Puccio, T., Kunka, K.S., An, S.S., and Kitten, T. (2022). Contribution of a ZIP-family protein to manganese uptake and infective endocarditis virulence in Streptococcus sanguinis. Mol Microbiol 117, 353–374.

26. Martini, A.M., Moricz, B.S., Ripperger, A.K., Tran, P.M., Sharp, M.E., Forsythe, A.N., Kulhankova, K., Salgado-Pabón, W., and Jones, B.D. (2020). Association of novel Streptococcus sanguinis virulence factors with pathogenesis in a native valve infective endocarditis model. Front Microbiol 11, 10.

27. Martini, A.M., Moricz, B.S., Woods, L.J., and Jones, B.D. (2021). Type IV Pili of Streptococcus sanguinis Contribute to Pathogenesis in Experimental Infective Endocarditis. Microbiol Spectr 9, e0175221.

28. Turner, L.S., Kanamoto, T., Unoki, T., Munro, C.L., Wu, H., and Kitten, T. (2009). Comprehensive evaluation of Streptococcus sanguinis cell wall-anchored proteins in early infective endocarditis. Infect Immun 77, 4966–4975.

29. Mecsas, J. (2002). Use of signature-tagged mutagenesis in pathogenesis studies. Curr Opin Microbiol 5, 33–37.

30. Xu, P., Ge, X., Chen, L., Wang, X., Dou, Y., Xu, J.Z., Patel, J.R., Stone, V., Trinh, M., Evans, K., et al. (2011). Genome-wide essential gene identification in *Streptococcus sanguinis*. Scientific reports 1, 125.

31. Zhu, B., Green, S.P., Ge, X., Puccio, T., Nadhem, H., Ge, H., Bao, L., Kitten, T., and Xu, P. (2021). Genome-wide identification of Streptococcus sanguinis fitness genes in human serum and discovery of potential selective drug targets. Mol Microbiol 115, 658–671.

32. Bao, L., Zhu, Z., Ismail, A., Zhu, B., Anandan, V., Whiteley, M., Kitten, T., and Xu, P. (2025). Experimental evolution of gene essentiality in bacteria. mBio 16, e0300525.

33. Abel, S., Abel zur Wiesch, P., Davis, B.M., and Waldor, M.K. (2015). Analysis of Bottlenecks in Experimental Models of Infection. PLoS Pathog 11, e1004823.

34. Senty Turner, L., Das, S., Kanamoto, T., Munro, C.L., and Kitten, T. (2009). Development of genetic tools for in vivo virulence analysis of Streptococcus sanguinis. Microbiology (Reading) 155, 2573–2582.

35. Gulsoy, I.C., Saaki, T.N.V., Wenzel, M., Syvertsson, S., Morimoto, T., Siersma, T.K., and Hamoen, L.W. (2025). Minimization of the Bacillus subtilis divisome suggests FtsZ and SepF can form an active Z-ring, and reveals the amino acid transporter BraB as a new cell division influencing factor. PLoS Genet 21, e1011567.

36. Puccio, T., An, S.S., Schultz, A.C., Lizarraga, C.A., Bryant, A.S., Culp, D.J., Burne, R.A., and Kitten, T. (2022). Manganese transport by Streptococcus sanguinis in acidic conditions and its impact on growth in vitro and in vivo. Mol Microbiol 117, 375–393.

37. Atkuri, K.R., Herzenberg, L.A., Niemi, A.K., and Cowan, T. (2007). Importance of culturing primary lymphocytes at physiological oxygen levels. Proc Natl Acad Sci U S A 104, 4547–4552.

38. Kawamura, Y., Hou, X.G., Sultana, F., Miura, H., and Ezaki, T. (1995). Determination of 16S rRNA sequences of Streptococcus mitis and Streptococcus gordonii and phylogenetic relationships among members of the genus Streptococcus. Int J Syst Bacteriol 45, 406–408.

39. Senthil Kumar, S., Johnson, M.D.L., and Wilson, J.E. (2024). Insights into the enigma of oral streptococci in carcinogenesis. Microbiol Mol Biol Rev 88, e0009523.

40. Abranches, J., Zeng, L., Kajfasz, J.K., Palmer, S.R., Chakraborty, B., Wen, Z.T., Richards, V.P., Brady, L.J., and Lemos, J.A. (2018). Biology of Oral Streptococci. Microbiol Spectr 6.

41. Hoshino, T., Fujiwara, T., and Kawabata, S. (2012). Evolution of cariogenic character in Streptococcus mutans: horizontal transmission of glycosyl hydrolase family 70 genes. Scientific reports 2, 518.

42. Shields, R.C., Zeng, L., Culp, D.J., and Burne, R.A. (2018). Genomewide identification of essential genes and fitness determinants of *Streptococcus mutans UA159*. mSphere 3.

43. Lambert, C., d’Orfani, A., Gaillard, M., Zhang, Q., Gloux, K., Poyart, C., and Fouet, A. (2023). Acyl-AcpB, a FabT corepressor in Streptococcus pyogenes. J Bacteriol 205, e0027423.

44. Ge, X., Kitten, T., Chen, Z., Lee, S.P., Munro, C.L., and Xu, P. (2008). Identification of Streptococcus sanguinis genes required for biofilm formation and examination of their role in endocarditis virulence. Infect Immun 76, 2551–2559.

45. Fan, J., Zhang, Y., Chuang-Smith, O.N., Frank, K.L., Guenther, B.D., Kern, M., Schlievert, P.M., and Herzberg, M.C. (2012). Ecto-5’-nucleotidase: a candidate virulence factor in Streptococcus sanguinis experimental endocarditis. PLoS One 7, e38059.

46. Morris, J.J., Lenski, R.E., and Zinser, E.R. (2012). The Black Queen Hypothesis: evolution of dependencies through adaptive gene loss. mBio 3.

47. Brilisauer, K., Rapp, J., Rath, P., Schöllhorn, A., Bleul, L., Weiß, E., Stahl, M., Grond, S., and Forchhammer, K. (2019). Cyanobacterial antimetabolite 7-deoxy-sedoheptulose blocks the shikimate pathway to inhibit the growth of prototrophic organisms. Nat Commun 10, 545.

48. Mir, R., Jallu, S., and Singh, T.P. (2015). The shikimate pathway: review of amino acid sequence, function and three-dimensional structures of the enzymes. Crit Rev Microbiol 41, 172–189.

49. Schönbrunn, E., Eschenburg, S., Shuttleworth, W.A., Schloss, J.V., Amrhein, N., Evans, J.N., and Kabsch, W. (2001). Interaction of the herbicide glyphosate with its target enzyme 5-enolpyruvylshikimate 3-phosphate synthase in atomic detail. Proc Natl Acad Sci U S A 98, 1376–1380.

50. Chakrabarti, M., Kannan, D., Munjal, A., Choudhary, H.H., Mishra, S., and Singh, S. (2020). Chorismate synthase mediates cerebral malaria pathogenesis by eliciting salicylic acid-dependent autophagy response in parasite. Biol Open 9.

51. Shende, V.V., Bauman, K.D., and Moore, B.S. (2024). The shikimate pathway: gateway to metabolic diversity. Nat Prod Rep 41, 604–648.

52. Spry, C., Kirk, K., and Saliba, K.J. (2008). Coenzyme A biosynthesis: an antimicrobial drug target. FEMS Microbiol Rev 32, 56–106.

53. Finkenwirth, F., and Eitinger, T. (2019). ECF-type ABC transporters for uptake of vitamins and transition metal ions into prokaryotic cells. Res Microbiol 170, 358–365.

54. Diamanti, E., Souza, P.C.T., Setyawati, I., Bousis, S., Monjas, L., Swier, L.J.Y.M., Shams, A., Tsarenko, A., Stanek, W.K., Jäger, M., et al. (2023). Identification of inhibitors targeting the energy-coupling factor (ECF) transporters. Commun Biol 6, 1182.

55. Guérin, H., Kulakauskas, S., and Chapot-Chartier, M.P. (2022). Structural variations and roles of rhamnose-rich cell wall polysaccharides in Gram-positive bacteria. The Journal of biological chemistry 298, 102488.

56. Martin, M. (2011). Cutadapt removes adapter sequences from high-throughput sequencing reads. EMBnet. journal 17, 10–11.

57. Langmead, B., and Salzberg, S.L. (2012). Fast gapped-read alignment with Bowtie 2. Nat Methods 9, 357–359.

58. Liao, Y., Smyth, G.K., and Shi, W. (2014). featureCounts: an efficient general purpose program for assigning sequence reads to genomic features. Bioinformatics 30, 923–930.

